# Comparative analysis of molecular signatures suggests the use of gabapentin for the management of endometriosis-associated pain

**DOI:** 10.1101/251397

**Authors:** Brice Bellessort, Anne Bachelot, Virginie Grouthier, Camille de Lombares, Nicolas Narboux-Neme, Paolo Garagnani, Chiara Pirazzini, Simonetta Astigiano, Luca Mastracci, Anastasia Fontaine, Gladys Alfama, Evelyne Duvernois-Berthet, Giovanni Levi

## Abstract

**Capsule:** Comparative analysis of gene expression signatures from endometriosis and mouse models shows that CACNAα2δs calcium-channel components involved in nociception are targets for the treatment of endometriosis-associated pain.

**Context:** Differential gene expression analyses comparing endometriotic lesions to eutopic endometrium have shown that the transcription factors *DLX5* and *DLX6* are drastically down-regulated in the ectopic implants. These finding suggests that regulatory cascades involving DLX5/6 might be involved in the origin of endometriosis symptoms such as chronic pelvic pain. We have shown that mice in which *Dlx5* and *Dlx6* are selectively inactivated in the uterus present an endometrial phenotype reminiscent of endometriosis implants.

**Objective:** Identify new targets for the treatment of endometriosis.

**Design:** To better focus the search for endometriosis targets we have compared the profile of genes deregulated in normal and ectopic women endometrium to those deregulated in the uterus of normal and *Dlx5/Dlx6*-null mice.

**Settings:** Academic research unit and University Hospital research laboratory

**Animals:** Mice carrying a uterus-specific deletion of *Dlx5/Dlx6*.

**Interventions:** Analysis of archive sections from normal endometrium and endometriosis implants.

**Main Outcome:** A novel endometriosis signature suggests that α2δs subunits of voltage-gated calcium channel are targets for the management of endometriosis-associated pain.

**Results:** We identify a signature of 30 genes similarly deregulated in human endometriosis implants and in *Dlx5/6*-null mouse uteri reinforcing the notion that the down-regulation of *Dlx5/6* is an early event in the progress of endometriosis. *CACNA2D3*, a component of the voltage-dependent calcium channel complex is strongly overexpressed both in endometriosis implants and in mutant mouse uteri; other members of the alfa2delta family, CACNA2D1 and CACNA2D2, are also overexpressed in endometriosis.

**Conclusion:** CACNA2D1, CACNA2D2 and CACNA2D3 are directly involved in pain perception. In particular, CACNA2D3 has been associated to pain sensitization and heat nociception in animal models while, in patients, variants of this gene are associated to reduced sensitivity to acute noxious stimuli. As CACNA2Ds are targets of gabapentinoids analgesics, our results suggest to consider the use of these drugs for the treatment of endometriosis-associated pain. Indeed, recent small-scale clinical studies have shown that gabapentin can be effective in the treatment of women chronic pelvic pain. Our findings reinforce the need for a large definitive trial.

## INTRODUCTION

Endometriosis is characterized by growth of endometrial-like tissue consisting of glandular tissue and stroma outside the uterine cavity in localized and innervated implants. Remarkably, endometriosis lesions present histopathological and physiological responses that are similar to those of the endometrium. Endometriosis patients frequently experience a variety of pain symptoms including chronic pelvic pain (CPP), dysmenorrhea, dyspareunia and dyschezia (1) that often result in severe personal, social and economic difficulties. Treatment of endometriosis-associated pain is difficult due to its poorly understood pathophysiology, its complex clinical presentation and natural history. Endometriosis is generally treated by either surgical intervention, by analgesic treatment or by hormonal treatment suppressing cyclic ovarian hormone production and reducing or eliminating menses (2). Current therapies present, however, limited efficacy on endometriosis pain, high rates of symptom recurrence, and significant side-effects (3, 4).

*Dlx* genes comprise a highly-conserved family of homeobox genes. Besides early roles in limb and craniofacial patterning during embryogenesis (5), *Dlx5* and *Dlx6* (*Dlx5/6*), are also involved in the differentiation of steroidogenic tissues including Leydig cells in the testis (6) and theca and granulosa cells in the ovary (7). *DLX5* and *DLX6* are expressed in glandular and lining epithelial cells of mouse and human endometrium (8) where their expression is upregulated during the secretory phase (9) of the cycle. By targeted inactivation of *Dlx5* and *Dlx6* in the mouse uterus, we have demonstrated their central role for uterine adenogenesis and fertility (8).

It has been repeatedly shown that the expression of *DLX5* and *DLX6* is strongly downregulated in endometriotic lesions compared to eutopic endometrium (10-13). Endometriosis implants and Dlx5/6 mutant mouse uteri have, therefore, in common a very low level of *Dlx5* expression. Furthermore, Dlx5/6-null endometrial epithelia present an histological appearance reminiscent of that found in endometriosis implants with very few glands and a cuboidal lining epithelium that locally tends to desquamate (8). The histological and molecular similarities between endometriosis implants and *Dlx5/6*-null mouse uteri have prompted our suggestion that the downregulation of *DLX5/6* could be an early event in the progress of endometriosis and that inactivation of these genes in the mouse uterus could constitute a model of the human pathology (8).

Here, we show that *Dlx5/6*-null mouse uteri and ectopic human endometrium share 30 deregulated genes compared to normal mouse uterus and eutopic endometrium confirming their similarity and permitting to focus existing endometriosis molecular signature. We focus our attention on *CACNA2D3* an over-expressed gene that has an evolutionary-conserved role in nociception (14). *CACNA2D1*, *CACNA2D2* and *CACNA2D3*, constitute the *α2δ* gene family of glycosylphosphatidylinositol-anchored (GPI-anchored) transmembrane voltage-gated calcium channels (VGCC)-associated subunits. During membrane depolarization, VGCCs permit Ca^2^ influx, contributing to cell depolarization and functioning as secondary messengers regulating key neuronal mechanisms including gene expression and neurotransmitter release. VGCCs contribute to the origin neuropathic pain and are targets of gabapentinoids (15). Remarkably, *CACNA2D1* to *3* are all significantly increased in endometriosis implants (11, 12) and should therefore be considered as targets for the treatment of endometriotic pain.

## MATERIALS AND METHODS

### Endometrial Biopsies

A total of 37 ovarian endometriosis and 12 normal endometrium samples were selected from the formalin-fixed, paraffin embedded tissue collection of the Division of Anatomic Pathology, Department of Surgical Science and Integrated Diagnostics (DISC) - University of Genoa, Italy. All patients signed an informed consent under local IRB approved protocols for research purposes dealing with the archived endometriosis tissues in the pathology department.

### Animals

Procedures involving animals were conducted in accordance with the directives of the European Community (council directive 86/609) and the French Agriculture Ministry (council directive 87–848, 19 October 1987, permissions 00782 to GL). Mice were housed in specific pathogen-free and light, temperature (21°C) and humidity (50-60% relative humidity) controlled conditions. Food and water were available ad libitum. Conditional *Dlx5/6*^*flox/flox*^ mutants were generated as described (8). To obtain *Pgr*^*cre*/+^; *Dlx5/6*^*flox/flox*^ mice, we crossed *Pgr*^*cre*/+^; *Dlx5/6*^*flox/+*^ males with *Dlx5/6*^*flox/flox*^ females.

### Immunohistochemistry

Fixed tissue samples were embedded in paraffin, and 5-mm thick sections were cut. Slides were dewaxed in xylol and re-hydrated in ethanol solutions. Antigen exposure was achieved submerging slides in sodium citrate 0.01 M, pH 6.0 (Sigma), and heating at 90 °C with pressure in a Retriever 2100 device (Prestige Medical). Endogenous peroxidase activity was quenched by incubation with 3% (vol/vol) H_2_O_2_ in methanol (Merck) or with peroxidase blocking solution from Dako Envision+System HRP kit (K4011, Agilent Pathology solutions, USA) for 30 minutes. Tissue sections were then blocked with 5% (wt/vol) bovine serum albumin (BSA) (Sigma) or 10% of Fetal Calf Serum (FCS) in PBS at room temperature for 2 hours, and then incubated with anti-DLX5 (NBP1-19547; Novus Biologicals, Littleton CO) polyclonal rabbit primary antibody at 1/200 dilution, or anti-CACNA2D3 at 1/250 dilution (NBP1-30557, Novus Biologicals, USA), or anti-CBS at 1/400 dilution (1478-1-AP, Proteintech PTGLAB, UK), or anti-LTBP2 at 1/250 dilution (17708-1-AP, Proteintech PTGLAB, UK), or anti-GJB3 at 1/50 dilution (12880-1-AP, Proteintech PTGLAB, UK), or anti-NOXA1 at 1/100 dilution (CUSABIO, USA), or anti-KIAA1324 at 1/200 dilution (ThermoFisher scientific, USA) in PBS 5% FCS overnight at 4°C in a humidified chamber. Normal rabbit IgGs were used as negative control. Next, slides were washed with PBS, and detection of the primary antibody was achieved by using the Histostain-SP Broad Spectrum kit (Invitrogen) or Goat anti-rabbit antibody linked to Horseradish peroxidase (Dako Envision+ System HRP kit; K4011, Agilent Pathology Solutions, USA). Counter-stain was done using Haematoxylin stabilized solution 1/5 (RAL diagnostics) according to standard procedures. Images were visualized on an Olympus BX-51 microscope and captured with Image-Pro Plus v.6.2 software (Media Cybernetics).

### RNA-seq

RNA library preparation was carried out according to the Illumina TruSeq Stranded Total RNA Sample Prep protocol, following Ribo-Zero Gold Deplete procedure (Illumina, San Diego, CA). From a total amount of 1.5 µg of total RNA, both cytoplasmic and mitochondrial ribosomal RNA (rRNA) were removed using specific biotinylated oligos and Ribo-Zero rRNA removal beads. RNA, purified from uteri of 3 *Pgr*^*Cre/+*^; *Dlx5/6*^*flox/flox*^ and 3 *Dlx5/6*^*flox/flox*^ mice at PN25, was fragmented by the addition of divalent cations and the incubation at 94°C for 4 minutes. The first strand cDNA was synthesized using random primers and a reverse transcriptase. DNA Polymerase I synthesized the second cDNA strand and generated blunt-ended ds cDNA, while RNase H removed the RNA template. An A nucleotide was added to the 3’ ends, followed by the ligation of multiple indexing adapters that are required for the pooling and the hybridization with the flow cell. These DNA products were purified and only those that have adapter molecules on both ends were enriched with PCR by using a Cocktail primer that anneals specifically to the adapters. The obtained libraries were validated by Bioanalyzer (Agilent DNA 1000, Agilent Technologies), pooled and sequenced using the Illumina HiSeq 2000 sequencing system. The global quality of each library was checked using FastQC software (http://www.bioinformatics.babraham.ac.uk/projects/fastqc/). To reduce any PCR amplification bias, choice has been done to remove duplicated reads before the mapping based on a 100% sequence identity between reads. Then, 5 nucleotides were trimmed at both 5’ and 3’ ends of all remnant reads using FASTX-Toolkit v0.0.14 (http://hannonlab.cshl.edu/fastx_toolkit/index.html). Reads were mapped against the mouse mm10 genome (version GRCm38.p3) using TOPHAT2 software v2.0.10 (16) with default parameters. Only uniquely mapped reads were kept and were counted on genes exons using HTSeq-count software v0.6.1p1 (17) with “union” mode using the *Mus musculus* Ensembl annotation (release 79). Only coding and expressed genes were considered. “R” package DESeq2 v1.8.0 (18) was used for statistical analyses to determine differential gene expression levels (cutoff: adjusted p-value≤0.1). Samples homogeneity was checked realizing a Principal Component Analysis with “R” package FactoMineR, following which, one mutant mouse was not considered for further analysis.

To identify mouse and human orthologs we used the Biomart tool, provided by Ensembl. From 424 genes differentially expressed in *Pgr*^*cre*/+^; *Dlx5/6*^*flox/flox*^ mice, we found 391 human orthologous which have been used for comparison.

### qPCR

Total RNA was isolated from five mouse uterine samples per group using an RNeasy minikit (Qiagen) according the manufacturer instructions. Deoxyribonuclease I (Roche) digestion was incorporated into an RNA isolation procedure to remove potential genomic DNA contamination. RNA concentration and the ratio of the absorbance at 260 and 280 nm were measured using a NanoDrop 2000 spectrophotometer (Thermo Scientific). Reverse transcription was carried out using 600 or 200 ng total RNA and Primscript (Ozyme) reverse transcriptase to obtain cDNA. Quantitative real-time PCR (qPCR) was performed using QuantStudio 6 Flex Real-time PCR System (Applied Biosystem). The PCR program consisted of 95°C for 10 min, 40 cycles 95°C of 15 seconds, 60°C for 10 min. Relative gene expression were measured with SYBR Green mix gene expression assays (Thermo Fisher). Primers utilized are shown below. To measure the relative amount of PCR products, the Ct geometric mean of *Sdha* and *Hprt1* was used as a control gene, was subtracted from the Ct of genes of interest to derive ∆Ct. The ∆Ct of mutant animals was compared with ∆Ct of control animals and the difference was assigned as ∆∆Ct. The fold change between two samples was then calculated as 2^-∆∆Ct^.

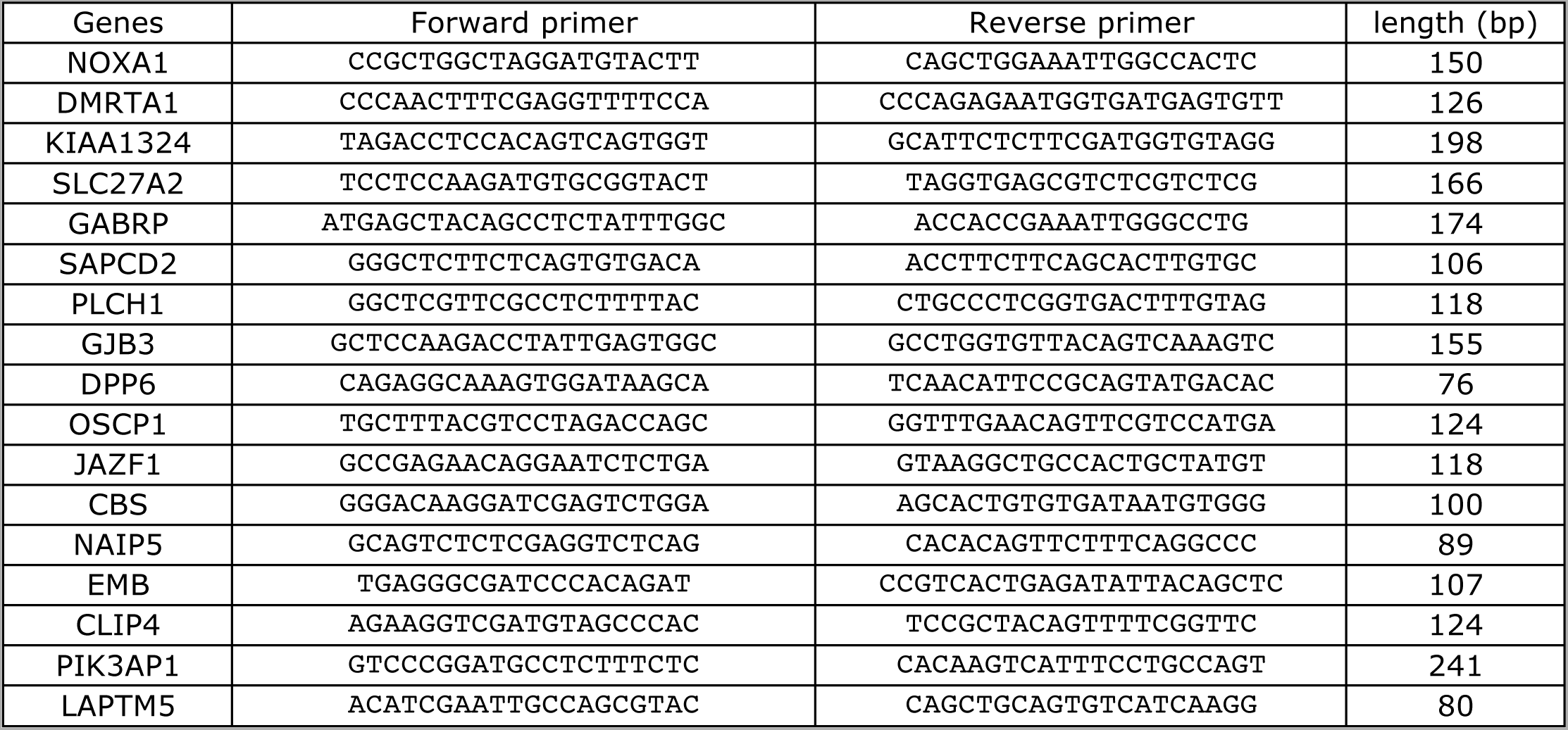

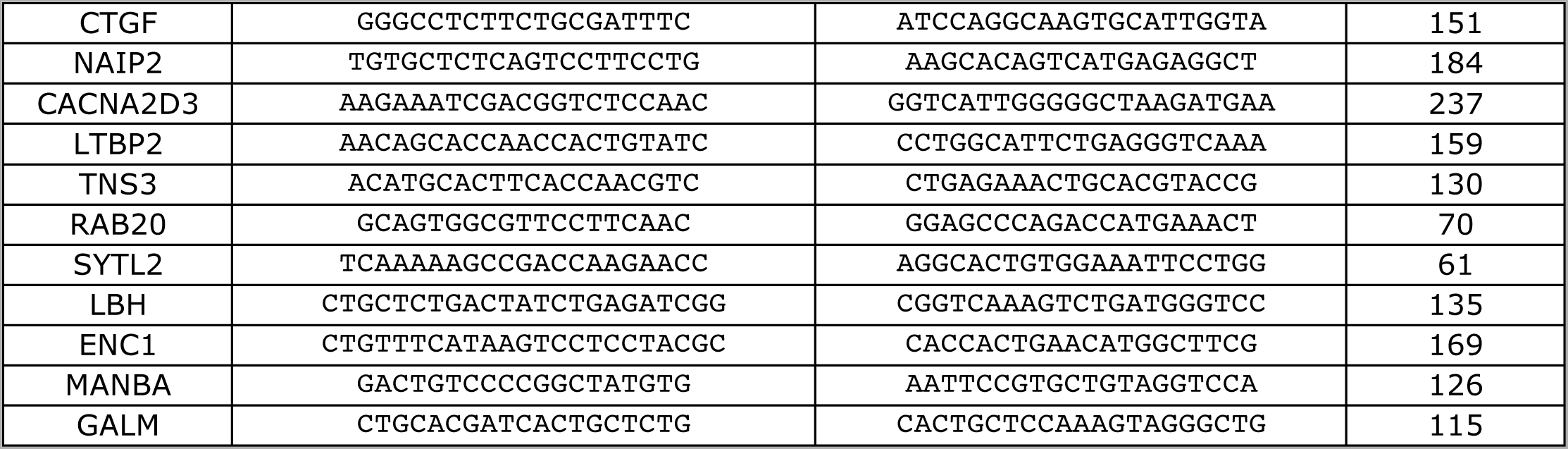

### Statistical analysis

The Mann-Whitney unpaired test was conducted using Prism (Graphpad Software, La Jolla, CA) to calculate the differences between groups. All values are expressed as means ± SEM of combined data from replicate experiments. Values of *P* < 0.05 were considered statistically significant.

## RESULTS

### DLX5 distribution in normal endometrium and in endometriosis

First, we analysed by immunohistochemistry the distribution of DLX5 in normal endometrium from various stages of the menstrual cycle (Fig.1A-B’). DLX5 protein is prevalently present in endometrial glandular and luminal epithelial cells. The intracellular distribution of DLX5 varies at different phases of the cycle: during the proliferative phase, DLX5 is detected both in the nucleus and in the cytoplasm (Fig.1A, A’), while, in the secretory phase, DLX5 is predominantly present in the cytoplasm (Fig.1B, B’) where it displays a polarized distribution mostly towards the luminal and basal aspects of the cells. Most endometriotic lesions (Fig.1C-D’) present very few glands with a very large lumen covered by an atypical non-columnar epithelium, which does not present a uniform phenotype even between different epithelial regions from the same patient (see for example the two facing epithelia in Fig. 1D’). In certain regions epithelial cells are large and non-cohesive (C’, D’) while in other territories they display a flattened phenotype with very little cytoplasm (D’, E’); cells are almost invariably ciliated. The intensity of anti-DLX5 staining is generally low or at the limit of detection (Fig. 1 C-D’), but, occasionally (Fig. 1E, E’), a clear signal can be detected in the cytoplasm and in perinuclear regions while nuclei are mostly negative; frequently epithelia are constituted by a mixture of DLX5-positive and DLX5-negative (arrows in E’) cells.

**Figure 1:**
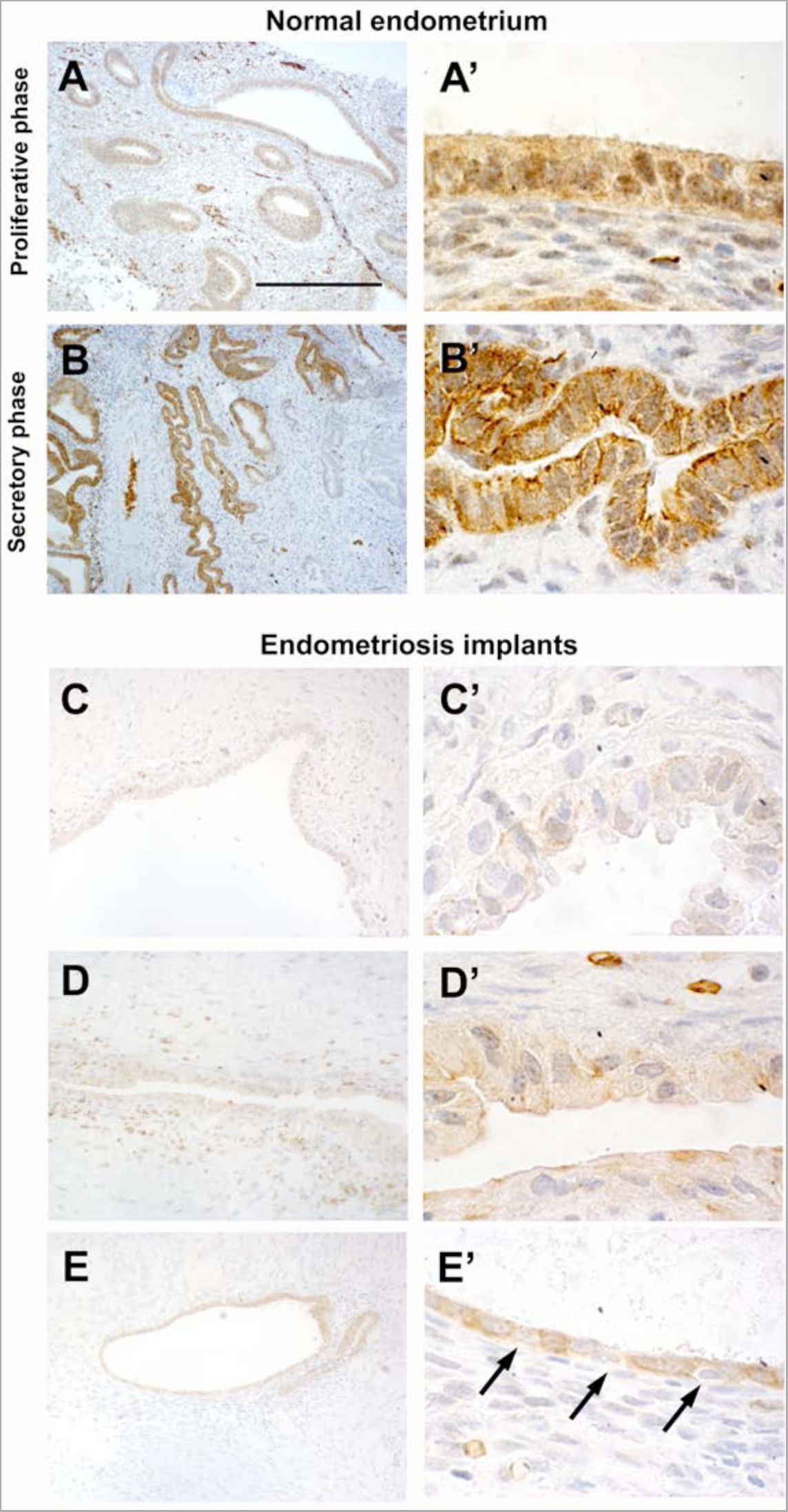
Immunohistlocalization of DLX5 in normal endometrium and ovarian endometriosis. In healthy endometrium, DLX5 protein is detected in glandular epithelia (A, A’, B, B’). During the proliferative phase (A, A’) the protein is present both in epithelial nuclei and in the cytoplasm. In contrast, during the secretory phase (B, B’) DLX5 presents a polarized distribution in the cytoplasm and is virtually absent from the nuclei. In ovarian endometriosis implants (C-E’) the expression of DLX5 is more variable. Most of the implants present very few glands with a very large lumen and a thin cuboidal epithelia which is in general constituted by a mixture of DLX5-positive and DLX5-negative (arrows in C’) cells. Scale bar: (A, B) 250 µm; (C, D, E) 125 µm; (A’, B’ C’, D’, E’) 25µm.

### A shared genetic signature between *Dlx5*/6-null mouse uteri and endometriosis implants

*DLX5* and *DLX6* are between the most down-regulated genes in endometriosis implants (10-12) and their inactivation in the mouse uterus results in an endometriosis-like phenotype (8). To better investigate the apparent similarity between endometriotic implants and mouse *Dlx5/6*-null uteri we have compared the RNAseq profiles of genes deregulated in the mutant mouse uterus to those reported in four independent studies in which the expression profile of endometriosis implants was compared to eutopic endometrium (11-13, 19). Hawkins *et al.* identified 2083 deregulated genes in endometriotic lesions (described on Supplementary table 2 of (11)); among this set of genes, 997 are under-expressed and 1086 are over-expressed. Several of these genes were also identified in the three other studies, which we have considered (12, 13, 19), providing a shared database to identify potential therapeutic targets. Comparison of *Dlx5/6*-mouse mutant uterus RNAseq dataset with the human datasets permits to identify 30 genes which present a common deregulation in the mouse and in endometriotic lesions: 13 down-regulated and 17 up-regulated (Table 1). If indeed the down-regulation of *Dlx5/6* is an early event in endometriosis progression, this set of 30 genes could well constitute a more focused molecular signature of endometriosis implants (Fig. 2).

**Table 1.** **Genes commonly deregulated after *Dlx5/6* inactivation in the mouse uterus and in human endometriosis implants** Genes presenting a common deregulation in *Pgr*^*cre*/+^; *Dlx5/6*^*flox/flox*^ mice and in studies comparing endometriosis implants to eutopic endometrium. In the mutant mouse *Dlx5* expression is absent

|  | Gene | Comparison of normal and *Dlx5/6*-null mouse uterus |  | Comparison of normal human endometrium and endometriosis implants |  |  |  |  |
| --- | --- | --- | --- | --- | --- | --- | --- | --- |
|  |  | log2 fold change | p value | log2 fold change Hawkins | p value Hawkins | Hever | Hull | Eyster |
| down | DLX5 | Absent (targeted deletion) |  | -8.434 | 1.804E-04 | down |  | down |
|  | NOXA1 | -2.25 | 3.51E-07 | -1.816 | 2.012E-03 | down |  |  |
|  | DMRTA1 | -2.11 | 2.65E-06 | -2.158 | 2.628E-04 |  |  |  |
|  | KIAA1324 | -2.07 | 1.96E-07 | -19.062 | 1.180E-05 | down | down | down |
|  | SLC27A2 | -1.91 | 1.14E-05 | -1.982 | 5.256E-03 | down | down |  |
|  | GABRP | -1.81 | 5.56E-06 | -22.036 | 2.671E-05 | down |  |  |
|  | SAPCD2 | -1.55 | 5.04E-04 | -2.155 | 6.115E-03 | down |  |  |
|  | CENPW | -1.36 | 6.92E-04 | -1.905 | 1.224E-04 | down |  |  |
|  | PLCH1 | -1.32 | 4.14E-04 | -3.082 | 5.696E-04 | down | down |  |
|  | GJB3 | -1.23 | 2.97E-03 | -1.835 | 5.448E-01 | down |  |  |
|  | DPP6 | -1.21 | 2.80E-03 | -2.917 | 2.170E-04 | down |  |  |
|  | OSCP1 | -0.95 | 3.24E-03 | -1.714 | 1.668E-03 |  |  | down |
|  | JAZF1 | -0.88 | 3.09E-03 | -2.719 | 2.028E-06 | down |  |  |
| up | CBS | 3.09 | 5.77E-27 | 2.491 | 3.844E-03 | up |  | up |
|  | NAIP5 | 2.44 | 1.69E-10 | 1.884 | 3.807E-04 |  |  |  |
|  | EMB | 1.76 | 4.38E-08 | 1.532 | 5.658E-03 |  |  |  |
|  | CLIP4 | 1.52 | 1.76E-08 | 2.612 | 2.198E-04 | up |  |  |
|  | PIK3AP1 | 1.48 | 8.44E-06 | 2.566 | 2.857E-03 | up |  |  |
|  | LAPTM5 | 1.45 | 8.85E-05 | 3.092 | 7.976E-03 | up |  | up |
|  | CTGF | 1.29 | 3.51E-04 | 1.582 | 7.149E-03 |  |  | up |
|  | NAIP2 | 1.19 | 1.04E-03 | 1.884 | 3.807E-04 |  |  |  |
|  | CACNA2D3 | 1.19 | 2.61E-03 | 2.066 | 2.469E-06 | up |  |  |
|  | LTBP2 | 1.14 | 3.16E-03 | 4.344 | 9.850E-05 | up | up | up |
|  | TNS3 | 1.12 | 1.22E-05 | 1.583 | 3.773E-03 | up | up |  |
|  | RAB20 | 1.08 | 3.26E-04 | 1.827 | 9.691E-03 |  |  |  |
|  | SYTL2 | 0.97 | 1.97E-03 | 1.788 | 8.826E-04 | up |  |  |
|  | LBH | 0.97 | 1.26E-03 | 2.029 | 1.615E-03 |  |  | up |
|  | ENC1 | 0.91 | 2.39E-03 | 2.919 | 3.286E-03 | up |  |  |
|  | MANBA | 0.83 | 1.98E-03 | 2.011 | 2.363E-03 | up |  |  |
|  | GALM | 0.76 | 2.27E-03 | 1.680 | 1.019E-03 | up |  |  |

**Figure 2:**
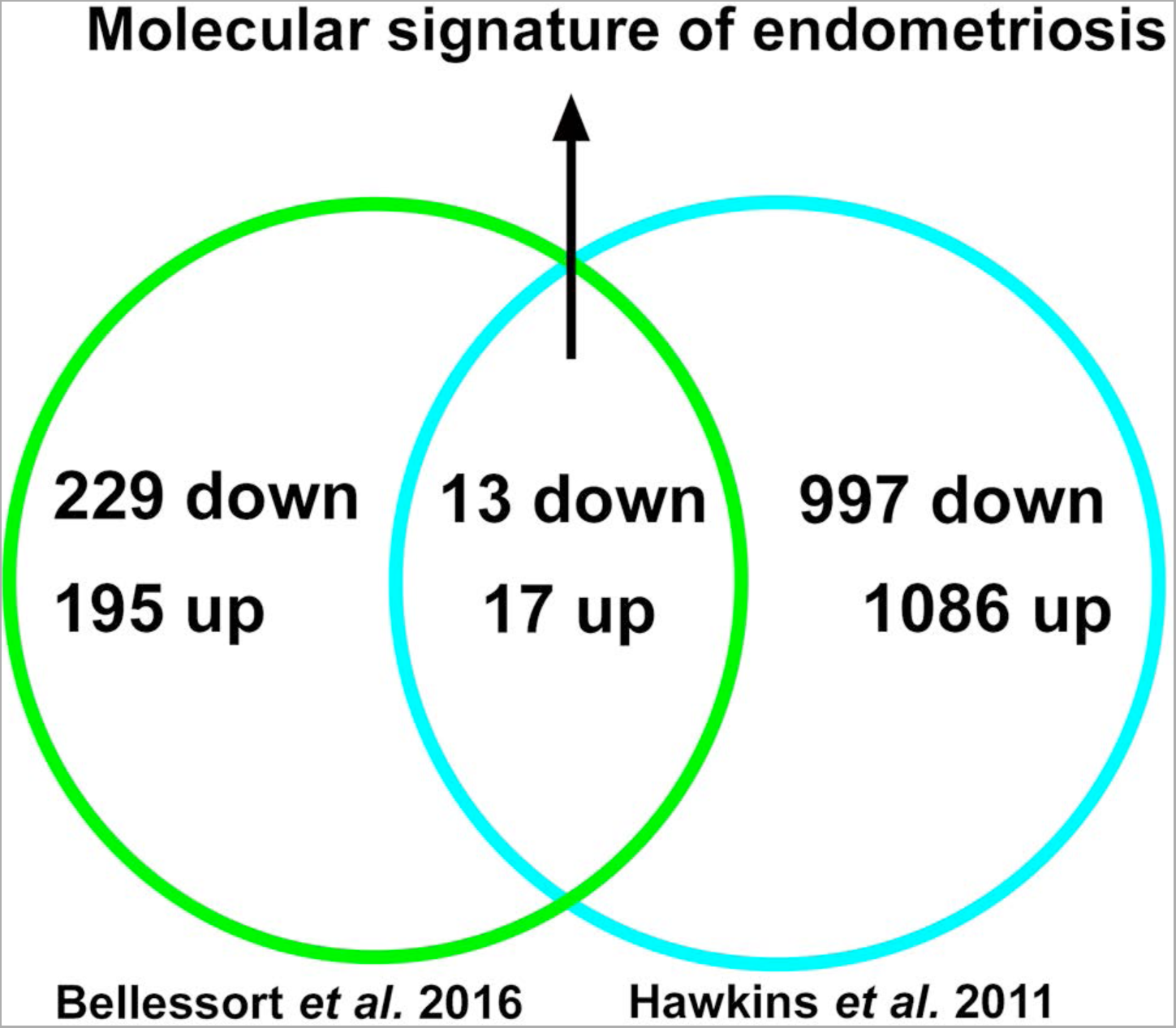
Molecular signature of endometriosis obtained comparing genes deregulated in *Dlx5/6* mutant mice uteri and in endometriosis lesions. The comparison between the Bellessort and Howkins datasets permits to propose a new molecular signature for endometriosis, see Table 1 for the names of individual genes.

### Validation by qPCR

In a first round of validation we analysed, by qPCR, the effect of *Dlx5/6* inactivation on the expression levels of 28 out of the 30 genes identified by RNAseq comparison (Figs. 3A,B). In all cases, the trend of variation occurred in the direction predicted by RNAseq either up (Fig. 3A) or down (Fig. 3B); however, due to high individual variability, the variation reached a level of significance for only 12 genes. Between the up-regulated genes, the expression of CACNA2D3 and CBS was respectively increased of 6 and 7 times. CACNA2D3 together with CACNA2D1 and CACNA2D2 constitutes the family of α2δ voltage gated calcium channel subunits involved in nociception. Remarkably, the expression of each of these three genes is significantly increased in endometriosis implants ((11, 12) and Data Set Records GDS2835, GDS3975). Interestengly, the Cystathionine-β-synthase (CBS) gene has been already shown to be hypomethylated and up regulated in endometriosis lesions (20).

**Figure 3:**
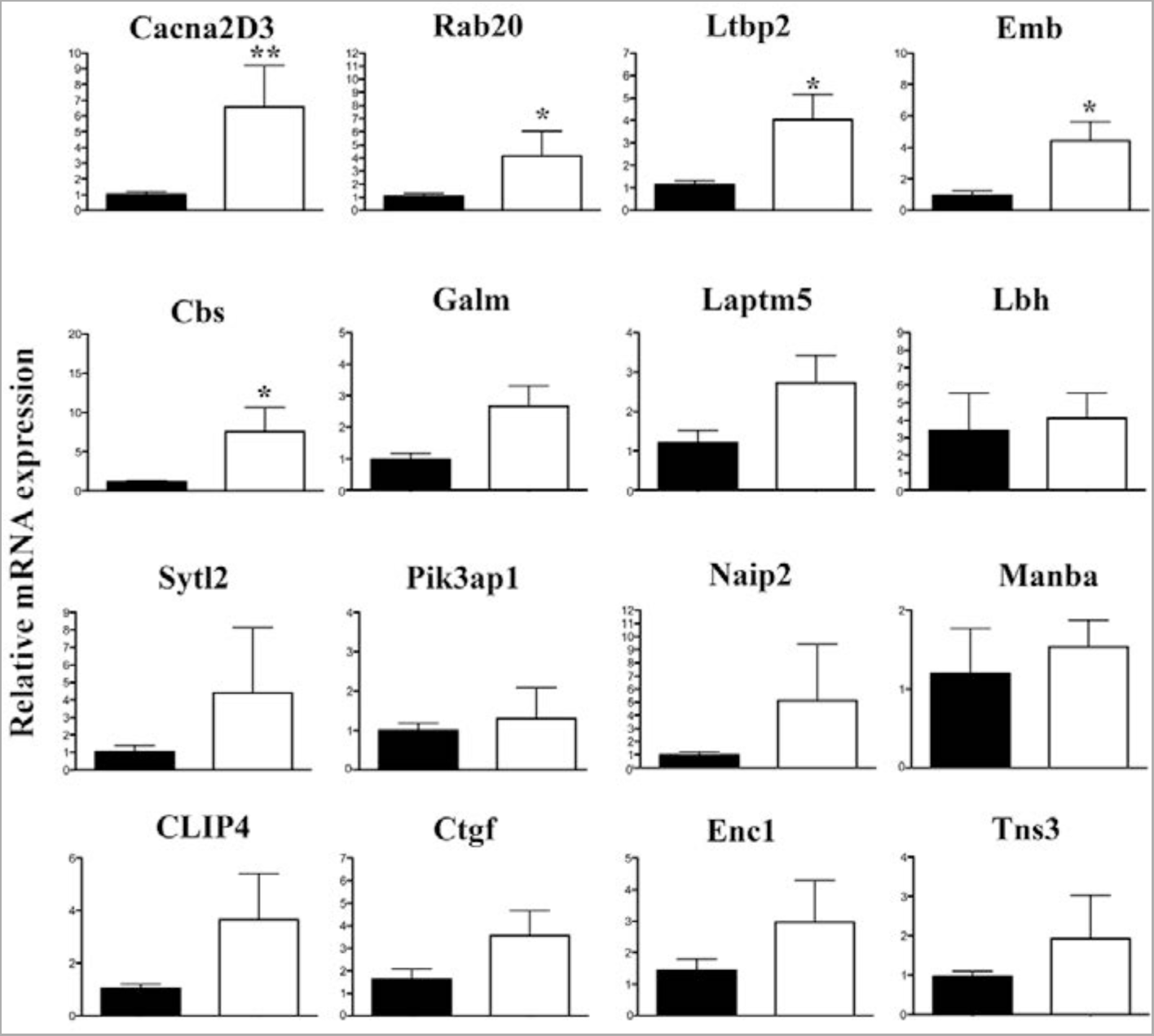
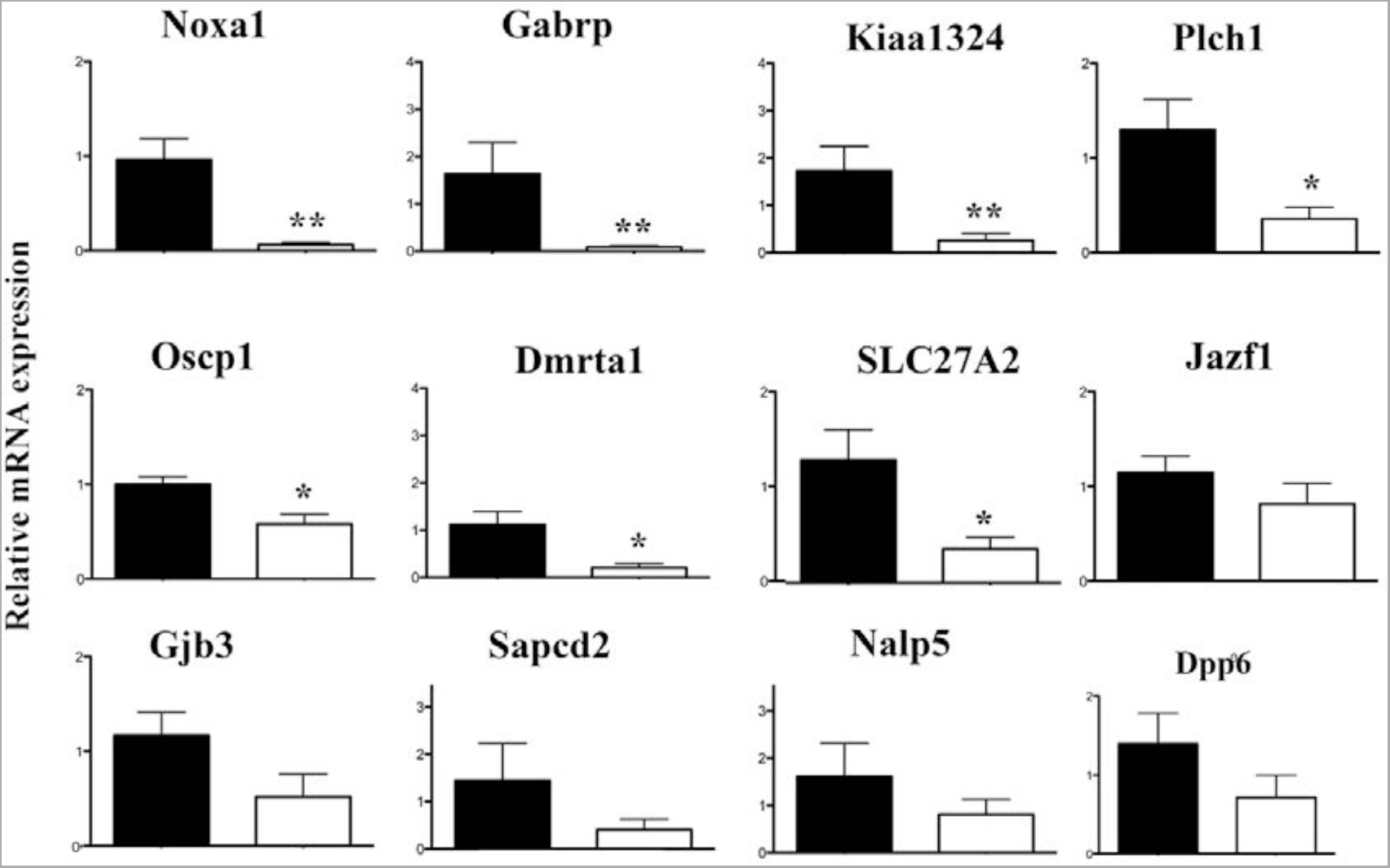
qPCR validation of genes upregulated after *Dlx5/6* invalidation in the mouse uterus. A) Relative mRNA expression in normal and mutant mouse uteri of upregulated genes identified as commonly deregulated in *Pgr*^*cre*/+^;*Dlx5/6*^*flox/flox*^ mice uteri and in endometriosis lesions;. Reference gene: *Sdha*. *= p<0,05, **=p<0,01, Mann and Whitney test analysis. B) Relative mRNA expression in normal and mutant mouse uteri of downregulated genes identified as commonly deregulated in *Pgr*^*cre*/+^;*Dlx5/6*^*flox/flox*^ mice uteri and in endometriosis lesions;. Reference gene: *Sdha*. *= p<0,05, **=p<0,01, Mann and Whitney test analysis.

### Altered pattern of gene expression in human endometriosis

To validate the up-regulation in endometriosis of the most significantly affected genes, we performed immunohistolocalization on section from normal endometrium and endometriosis implants. CACNA2D3, CBS and LTBP2 (Fig. 4) immunodetection signal was dramatically increased in endometriotic lesions. CACNA2D3 was virtually undetectable on normal endometrium sections (Fig. 4 A,B) and presented a high level of expression in endometriosis epithelia and smooth muscle cells (Fig. 4 A’,B’). In the epithelia, CACNA2D3 immunodetection was predominantly associated to membranes and cytoplasm (arrows in Fig. 4 B’) while CBS and LTBP2 immunodetection (Fig. 4 C-D’) was mostly nuclear in endometriosis implants.

**Figure 4:**
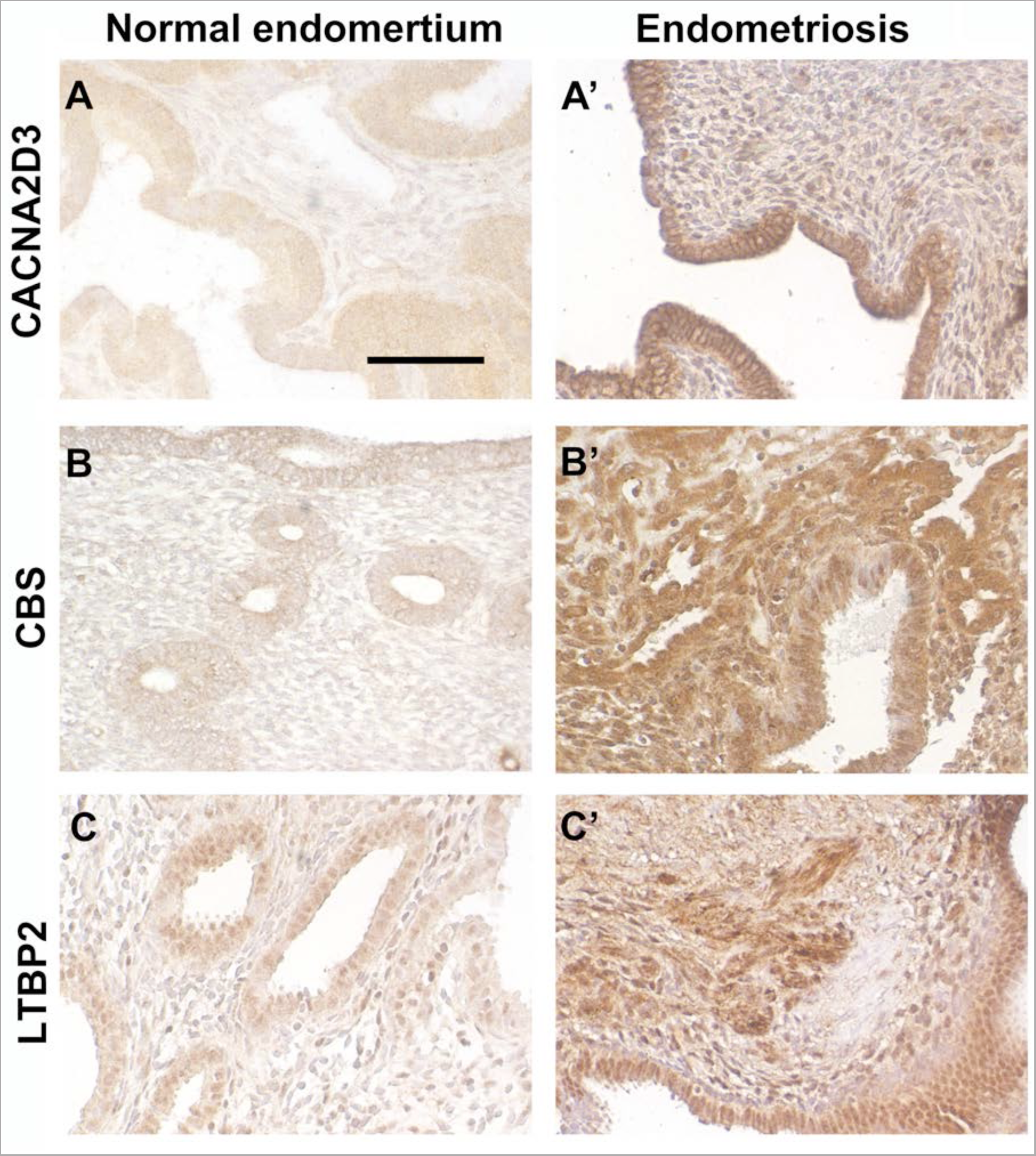
Immunolocalization of CACNA2D3, CBS and LTBP2 in normal endometrium and in ovarian endometriosis implants. Immunolocalization of CACNA2D3, CBS and LTBP2 on sections from normal endometrium (A, B, C) and from endometriosis implants (A’, B’, C’). Arrows indicate CACNA2D3 distribution in epithelial cells of endometriosis implants. Scale bar: (A, A’; C, C’, D, D’) 250 µm; (B, B’) 62,5 µm.

## DISSCUSSION

Morphological and genetic observations on mutant mouse uteri and on pathological specimens have prompted the suggestion that low *DLX5* and *DLX6* expression could be associated to the progression of endometriosis. We have therefore proposed that *Dlx5/6*-null mouse uteri could be considered as a model of the disease permitting to get new insight on this pathology (8).

Here we have compared the gene expression signature of *Dlx5/6*-null mouse uteri to those obtained in four independent studies on endometriosis (11-13, 19). *Dlx5/6*-null uteri and endometriosis implants share a set of 30 commonly deregulated genes supporting the hypothesis that down-regulation of *Dlx5/6* could be an early event in endometriosis progression. The 30 identified genes constitute a more “focused” genetic signature of the disease that has permitted to handpick potential therapeutic targets. We focused, in particular, on genes up-regulated in endometriosis as they could be potential targets for therapeutic strategies.

Current endometriosis therapies present limited efficacy, high rates of symptom recurrence, and significant side-effects (3, 4). For example the use of GnRH agonists is associated to side effects such as menopausal symptoms and reduced bone mineral density (21) which restrict their use and demand add-back therapy (22). In addition, pain management is one of the major clinical challenges for the treatment of endometriosis (23). Endometriosis implants, although ectopic, are highly innervated. Peritoneal endometriosis lesions present a high density of sensory, cholinergic and adrenergic nerve fibres that might be involved in the origin of pain (24, 25). It has been proposed that a bidirectional relation exists between nerve fibres and endometriotic lesions with a two-way interaction between the implants and the central nervous system (26-28). The interplay between the peripheral and the central nervous system might be at the origin of individual differences in pain perception that can, in some patients, evolve independently from the disease (29). When pathological conditions cause damage to the nervous system, sensory activation thresholds are lowered with exaggerated pain perception even upon mild or absent painful stimulation.

Between the 17 up-regulated gene candidates, we focused specifically on CACNA2D3, member of the family of α2δ voltage gated calcium channel (VGCC) subunits involved in nociception (Summary diagram in Fig. 5). CACNA2D1, CACNA2D2, CACNA2D3 and CACNA2D4 constitute the α2δ family of VGCC subunits (15). VGCC are composed of α1 subunit proteins involved in pore formation (for review see (30)) and many possible splice variants of the α1, β and α2δ subunits. Such a molecular diversity confers to these channels a variety of specific properties in different cell types and situations. VGCCs play a critical role in neuropathic pain development through the modulation of the release of excitatory neurotransmitters (31), calcium-dependent enzyme activation (32), gene regulation (32-34) and short-and long-term plasticity changes (35-38). Abnormal regulation of VGCC subunits, such as α2δs, may contribute to pain signal transduction through several mechanisms including the induction of abnormal synaptogenesis (39, 40). Dysregulation of VGCCs and their subunits has been observed in pathological conditions, including nerve injuries and animal models of endometriosis (41); it is therefore possible that these channels contribute to the origin of endometriosis-associated pain.

**Figure 5:**
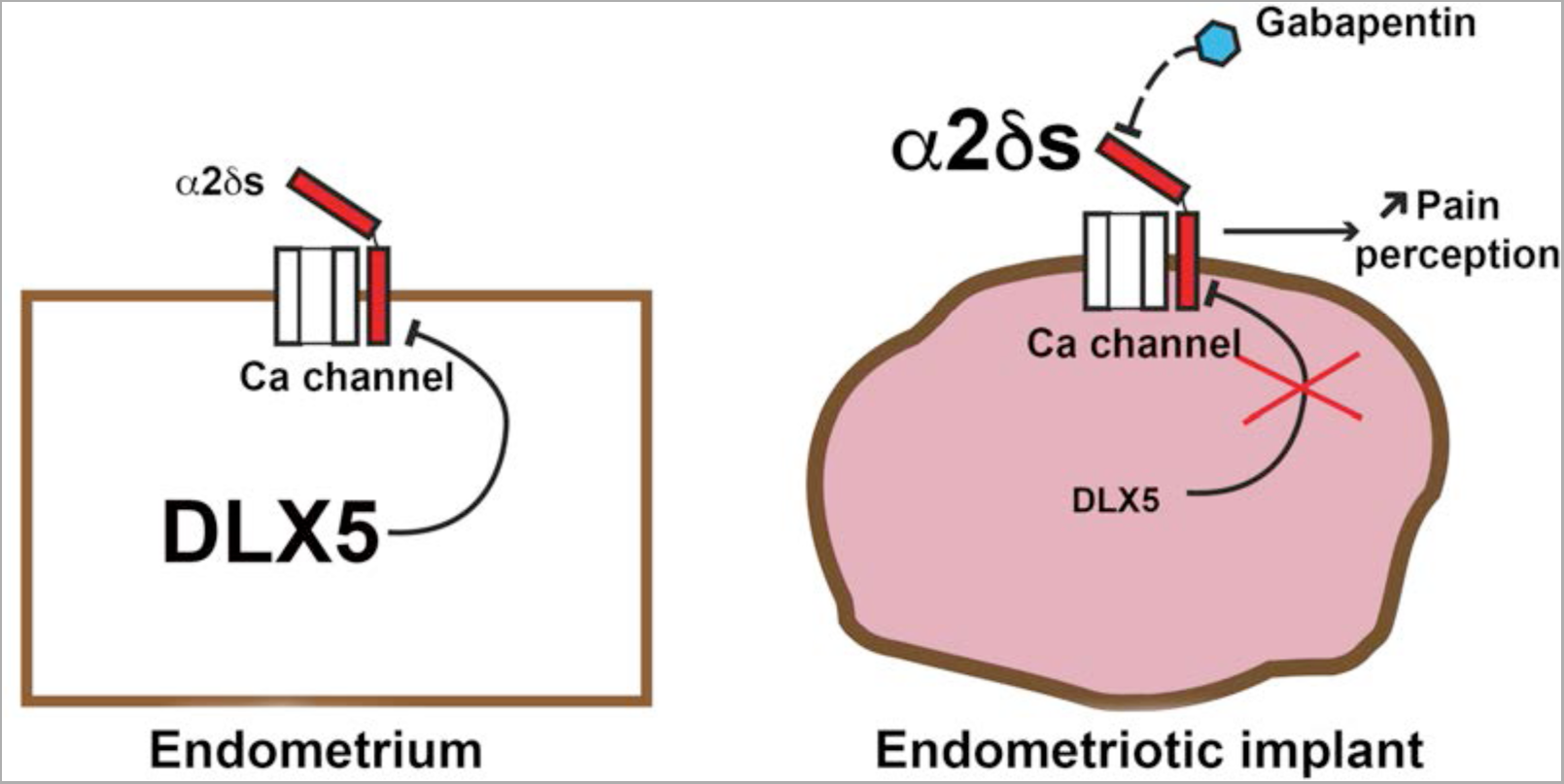
Summary diagram. The reduction of *DLX5* expression in endometriosis implants results in increased expression of *α2δ* subunits with a potential increase in neuropathic nociception.

Gabapentinoids, including gabapentin (Neurontin; Pfizer) and pregabalin (Lyrica; Pfizer), are compounds used for the treatment of neuropathic pain (42, 43). As seen by binding of [^3^H]Gabapentin to membranes from COS-7 cells transfected with α2δ cDNA, gabapentinoids bind predominantly to α2δ1, α2δ2 subunits of VGCCs (44-46). However, an evolutionary conserved role in nociception has been shown also for α2δ3 and SNP variants of this subunit are associated to reduced sensitivity to heat and chronic back pain. (14).

Analgesics, such as non-steroidal anti-inflammatory drugs (NSAIDs) or gabapentin are often used for pain relief despite limited evidence of their efficacy in endometriosis (4, 47). A recent pilot trial on 47 patients (48) and several clinical observations (see for example (47)) have suggested that treatment with gabapentin alleviates significantly endometriotic pain, however given the small size of the trials uncertainty remains (42). Our findings provide evidence supporting a definitive evaluation of the efficacy of gabapentin in the management of endometriosis-associated pain.

## ACKNOWLEDGEMENTS

We thank Dr Elisabeth Da Maia, Department of Pathology, Pitié Salpêtrière Hospital, for her kind help in endometrial and endometriosis section analysis.

## FUNDING

This study was supported in part by the French Centre National de la Recherche Scientifique, by the French National Museum of Natural History and by the EU Consortium HUMAN (EUFP7-HEALTH-602757). - The authors report no conflict of interest.

